# OME Files - An open source reference library for the OME-XML metadata model and the OME-TIFF file format

**DOI:** 10.1101/088740

**Authors:** Roger Leigh, David Gault, Melissa Linkert, Jean-Marie Burel, Josh Moore, Sébastien Besson, Jason R. Swedlow

## Abstract

Digital imaging is now used throughout biological and biomedical research to measure the architecture, composition and dynamics of cells, tissues and organisms. The many different imaging technologies create data in many different formats. This diversity of data formats arises because the creation of common, cross-domain data standards is difficult and the pace of innovation challenges the durability of any defined standard and the implementation of data stewardship principles such as FAIR^1^. The Open Microscopy Environment (OME; http://openmicroscopy.org) has therefore developed the OME Data Model, an open, extensible specification for imaging metadata, that supports metadata related to an imaging experiment, data acquisition and any derived analytic results^2–4^. The OME Data Model has been implemented in OME-XML so that metadata can be stored and accessed by any software. OME-TIFF embeds OME-XML into the header of a TIFF file, making scientific imaging metadata accessible to any software that can read the TIFF format. To support the usage of OME-XML and OME-TIFF, OME has built Bio-Formats, a Java-based library that converts the metadata in proprietary scientific image file formats into the open OME Data Model^5^.

Bio-Formats is used in Fiji, OMERO and many other applications to read metadata from scientific image file formats^6,7^, but because it is written in Java, it has not been included in software that controls most imaging systems. We have therefore ported all the image file input, output and metadata functions in Bio-Formats into a completely new C++-based project called OME Files. OME Files (http://www.openmicroscopy.org/site/products/ome-files-cpp) is freely available (https://github.com/ome/ome-files-cpp), liberally licensed (BSD-2) and builds on all major platforms (http://downloads.openmicroscopy.org/ome-files-cpp/), so it can be used by commercial and academic software projects. Example uses include image data acquisition, which may require low level access to the acquisition hardware and subsequent writing of the acquired data as OME-TIFF, and image analysis which may require the use of C++ libraries such as OpenCV, Eigen or ITK/VTK, using OME-TIFF as the file format for the input data and any transformed output data. In both of these examples, OME Files will read and write the image data and metadata using OME-XML and OME-TIFF to provide interoperability with the wide range of existing software supporting these open formats. The experimental and acquisition metadata may be stored as OME-XML metadata when writing at acquisition time, and read during subsequent analysis. Together OME Files and Bio-Formats provide solutions for image metadata access across the Java and C++ programming domains (Fig. 1).

**Figure 1.**
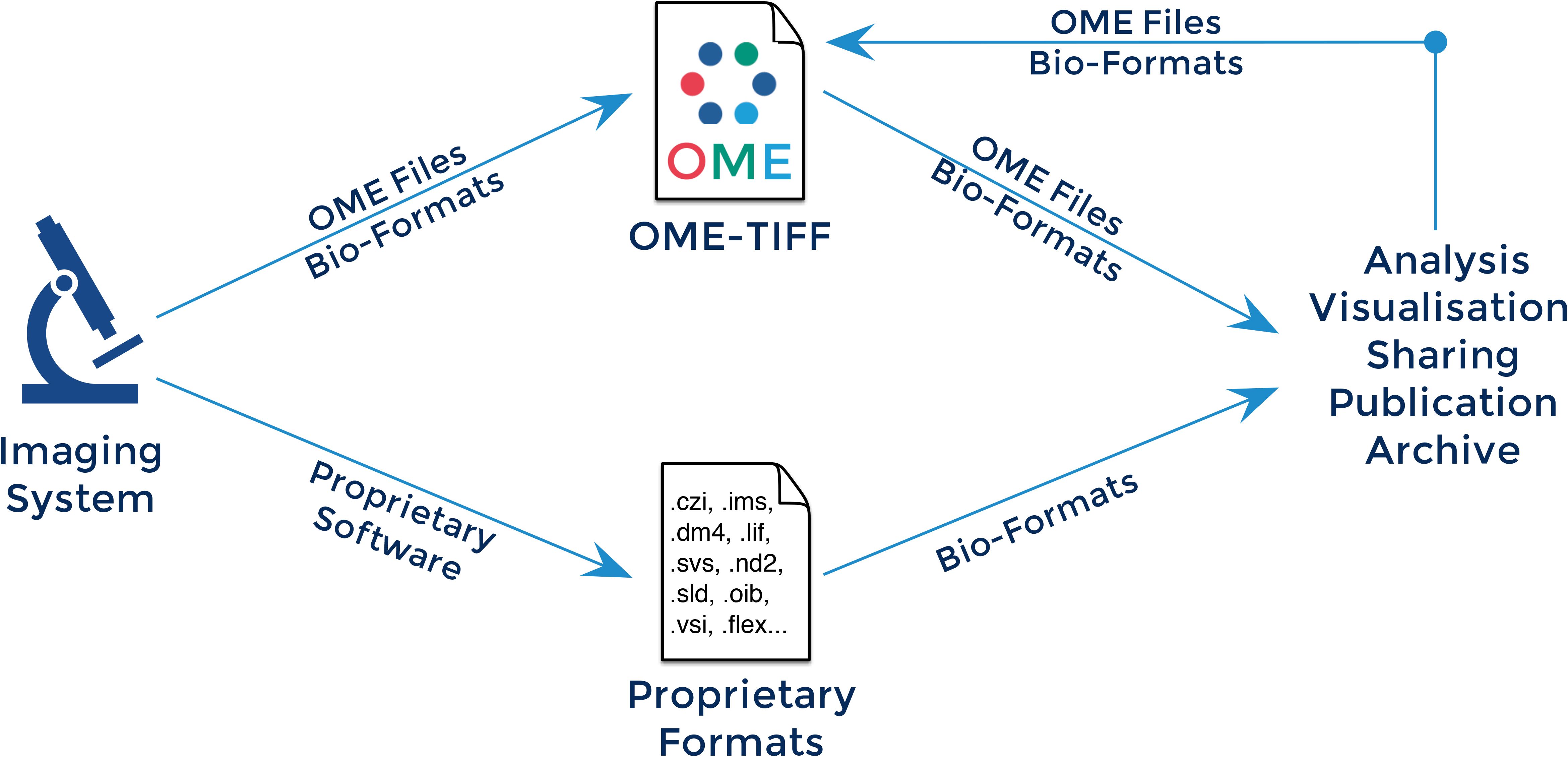
The image acquisition and analysis workflow, focussing on the software and formats that are used. Binary pixel data is combined with experimental and acquisition metadata and, the result stored in either proprietary or open OME-TIFF data files. Software using OME Files (C++) or Bio-Formats (Java) creates OME-TIFF directly. Alternatively, data are written in proprietary or custom formats. OME Files and Bio-Formats, used as libraries inside image processing and analysis software, will read and write OME-TIFF files, ensuring that the processing results are preserved in an open, accessible format. Bio-Formats can additionally read many proprietary formats, and serves as a way to ensure interoperability between closed formats and other software. It can also be used to convert proprietary formats into open, accessible formats such as OME-TIFF.

To test the utility of OME Files, we developed benchmark tests for Bio-Formats, OME Files and a Java Native Interface (JNI) library that is an alternative implementation that bridges Bio-Formats to C++^8^. For benchmark data, we converted a multi-dimensional fluorescence image, a high content screening plate (derived from the BBBC resource^9^), and a time-lapse sequence with >10,000 associated regions of interest (ROIs; derived from the MitoCheck study^10^) to OME-TIFF. For each of these benchmark files, we determined the relative performance for each library as the ratio of read and write times for image metadata and binary pixeldata using OME Files or JNI to the same times recorded using Bio-Formats on either Linux or Windows operating systems (Fig. 2; see Supplemental Information). We ran each test multiple times to account for any variations in system activity or other factors. Figure 2 shows that OME Files is ~10× or more faster than using Bio-Formats reading binary pixel data from all benchmarks, and is also much faster than Bio-Formats writing these data. This improved performance is important for handling large data volumes created by modern scientific cameras. OME Files is several-fold faster reading and writing metadata from a multi-dimensional fluorescence image. For large, complex metadata sets, OME Files is faster than Bio-Formats for writing metadata, but slower for reading metadata, although these differences are relatively small. The JNI implementation performs quite similarly to Bio-Formats for all reading and writing operations except for writing pixeldata where it is much slower than OME Files in all three benchmark tests. These performance metrics suggest that OME Files is a performant, useful resource for reading and writing OME-TIFF in several different use cases.

**Figure 2.**
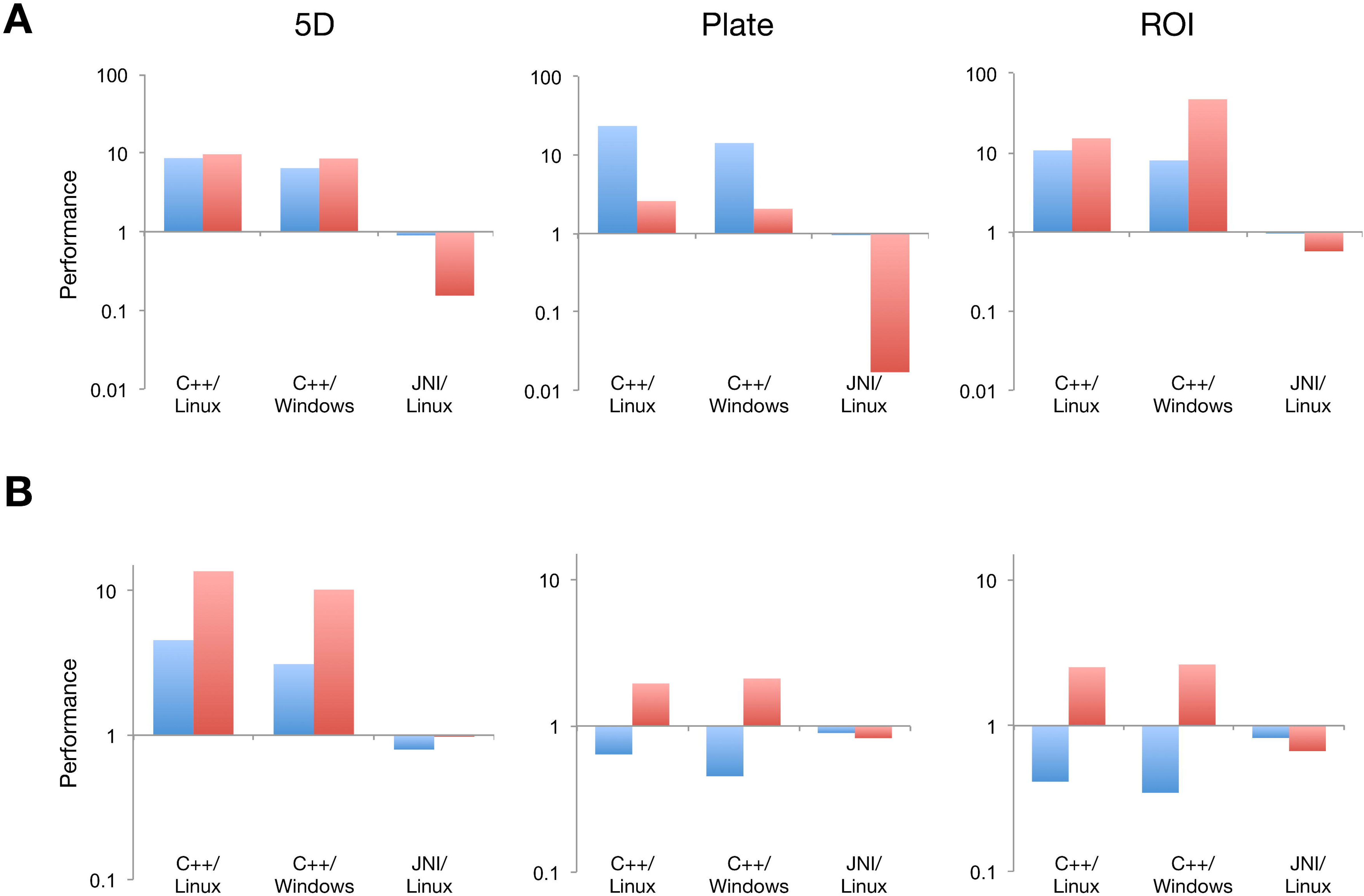
Performance comparison of the OME Files C++ and JNI libraries. The reading (blue) and writing (red) execution times for (A) pixeldata and (B) metadata of three OME-TIFF-based datasets were measured using the OME Files C++ library and the JNI implementation that embeds Bio-Formats within C++. For each benchmark test, we expressed performance as the inverse of the execution time and normalized it by the performance of Bio-Formats against the same dataset under the same operating system (i.e., Windows or Linux). We measured relative performance for three benchmark datasets, a multi-dimensional fluorescence image (5D), a high-content screening plate (Plate) and a time-lapse sequence with >10,000 associated ROIs. Standard deviations calculated from multiple repetitions were found to be systematically smaller than 9% and not included in the figure. For more information about the benchmark tests, datasets and the tabular results including standard deviations, see Supplementary Information and Supplementary Table 1.

OME-TIFF has been adopted by many commercial imaging software providers. However since commercial software is often written in native programming languages such as C++, each has implemented its own version of the OME-TIFF readers and writers. OME Files provides a way to minimise this divergence, as it is built and tested on all major compilers and operating systems. In addition, since large-scale calculations most often use native code, OME Files provides a seamless, performant way for these calculations to use OME-TIFF. Finally, OME Files makes it much easier to support OME-TIFF usage in emerging computational environments like iPython notebooks and to explore the development of next generation file formats using containers such as HDF5 to provide scalable access to large and complex image datasets, as well as to support *n*-dimensional datasets^11^.

## Acknowledgements

The authors thank the OME user community for helpful feedback and suggestions for improvements to OME software. This work was supported by a Wellcome Trust Strategic Award (095931/Z/11/Z) and two BBSRC BBR awards (BB/L024233/1 and BB/M018423/1).

